# Experimental evolution for improved post-infection survival selects for increased disease resistance in *Drosophila melanogaster*

**DOI:** 10.1101/2024.02.14.580293

**Authors:** Aabeer Basu, Kimaya Tekade, Aparajita Singh, Paresh Nath Das, Nagaraj Guru Prasad

## Abstract

Disease resistance (defined as the host capacity to limit systemic infection intensity) and disease tolerance (defined as the host capacity to limit infection-induced damage) are two complementary defense strategies that help the hosts maximize their survival and fitness when infected with pathogens and parasites. In addition to the underlying physiological mechanisms, existing theory postulates that these two strategies differ in terms of the conditions under which each strategy evolves in host populations, their evolutionary dynamics, and the ecological and epidemiological consequences of their evolution. Here we explored if one or both of these strategies evolve when host populations are subjected to selection for increased post-infection survival. We experimentally evolved *Drosophila melanogaster* populations, selecting for the flies that survived an infection with the entomopathogen *Enterococcus faecalis*, and found that the host populations evolved increased disease resistance in response. This was despite the physiological costs associated with increased resistance. We did not find evidence of any change in disease tolerance in the host populations. We have therefore demonstrated that in an experimental evolution set-up, where insect hosts must survive an infection with a pathogenic bacterium, the hosts evolve improved disease resistance but not disease tolerance.

## 1. Introduction

In order to survive a virulent systemic infection, the infected host must prevent proliferation of the invading pathogen and minimize systemic pathogen load. Simultaneously, the infected host must mitigate any damage to its system brought about due to the infection. Therefore, both disease resistance (i.e., the host ability to minimize systemic pathogen burden) and disease tolerance (i.e., the host ability to minimize infection induced damages) are the available alternate, and somewhat complementary, defense strategies an infected host utilizes to maximize its survival (Medzhitov et al., 2008, Read et al., 2008, Schneider and Ayres 2008, Raberg et al., 2009, Raberg and Stjernman 2012, Ayres and Schneider 2012, Raberg 2014, Kutzer and Armitage 2016, Lissner and Schneider 2018). The eco-evolutionary dynamics for these two host defense strategies are distinct from one another and has been elucidated in detail in various theoretical studies (Roy and Kirchner 2000, Restif and Koella 2004, Miller et al., 2006, Miller et al., 2007, Best et al., 2008, Carval and Ferriere 2010, Best et al., 2014, Singh and Best 2021).

The genetic architecture of both disease resistance and tolerance has also been investigated in various animal model systems (Hansen and Koella 2003, Raberg et al., 2007, Lefevre et al., 2011, Parker et al., 2014). In *Drosophila melanogaster*, genetic variation for both traits have been documented in both laboratory and wild populations, and single-gene mutations are known that can modify either or both defense strategies (Lazzaro et al., 2004, Lazzaro et al., 2006, Corby-Harris et al., 2007, Ayres and Schneider 2008, Ayres et al., 2008, Dionne and Schneider 2008, Howick and Lazzaro 2014, Vincent and Sharp 2014, Howick and Lazzaro 2017, Hotson and Schneider 2015, Duneau et al., 2017b, Miller and Cotter 2018). Additionally, environmental factors that can influence these strategies have also been identified (Lambrechts et al., 2006, Corby-Harris and Promislow 2008, Ayres and Schneider 2009, Zeller and Koella 2017, Cumnock et al., 2018). Despite these empirical investigations, and a variety of theoretical studies describing the eco-evolutionary dynamics of resistance and tolerance, demonstration of real-time evolution of either of these strategies – either in the wild (Behrman et al., 2017) or in the laboratory (Zeller and Koella 2017, Silva et al., 2021) – has been rare.

In the present study we explore if disease resistance or tolerance evolves when *Drosophila melanogaster* populations are subjected to long-term directional selection for increased post-infection survival in a laboratory experimental evolution set-up. Although resistance or tolerance can evolve in response to direct selection on these traits, such specific direct selection is unlikely in nature. In natural settings, hosts are under selection to maximize their post-infection fitness, and in such conditions either or both strategies can be selected for, since both strategies offer the solution to the problem of maximizing post-infection survival. The strategy which the hosts evolve will probably be dependent on the exact ecology of the host, and contingent on various other factors such as standing genetic variation for either trait in the host population, genetic correlation between these traits, existing levels of each trait, and the costs associated with their evolution, both physiological and ecological (Restif and Koella 2004, Boots et al., 2009, Raberg and Stjernman 2012, Best et al., 2014, Singh and Best 2021). It is therefore pertinent to test the evolution of disease resistance and/or tolerance in response to selection for increased post-infection survival.

We experimentally evolved replicate *Drosophila melanogaster* populations to better survive systemic infection with a Gram-positive bacterial pathogen, *Enterococcus faecalis*. Within 35 generations of forward selection, the selected populations exhibited significant increase in post-infection survival compared to the control populations (Singh et al., 2021). Thereafter, we tested if this increase in survival in the selected populations was due to increased resistance, increased tolerance, or both. The results suggest that the selected populations are better at restricting systemic pathogen proliferation after being infected but carry similar bacterial loads at the point of mortality, compared to the control populations. We propose that these observations suggest that the selected populations have evolved to become more resistant to infection, without any measurable change in their tolerance levels.

## 2. Material and methods

### 2.1. Host populations

Replicate *Drosophila melanogaster* populations were evolved parallelly, subjecting some of the populations to selection for increased post-infection survival, while maintaining the others as either procedural or uninfected controls. The experimental evolution set-up consisted of 12 populations in total, distributed into 3 selection regimes (**figure 1**, Singh et al., 2021):

A. **E_1-4_: Populations selected for better survival following infection with the Gram-positive bacterium *Enterococcus faecalis*.** Every generation, 2–3-days old adult flies (200 females and 200 males) are subjected to infection with *E. faecalis*, and 96-hours post-infection, the survivors are allowed to reproduce and contribute to the next generation. At the end of 96 hours, on average 100 females and 100 males are left alive in each of the E_1-4_ populations.
B. **P_1-4_: Procedural (sham-infected) control populations.** Every generation, 2–3-days old adult flies (100 females and 100 males) are subjected to sham-infection, and 96-hours post-sham-infection, the survivors are allowed to reproduce and contribute to the next generation.
C. **N_1-4_: Uninfected control populations.** Every generation, 2–3-days old adult flies (100 females and 100 males) are subjected to light CO_2_ anesthesia only, and 96-hours post-procedure, the survivors are allowed to reproduce and contribute to the next generation.

**Figure 1.**
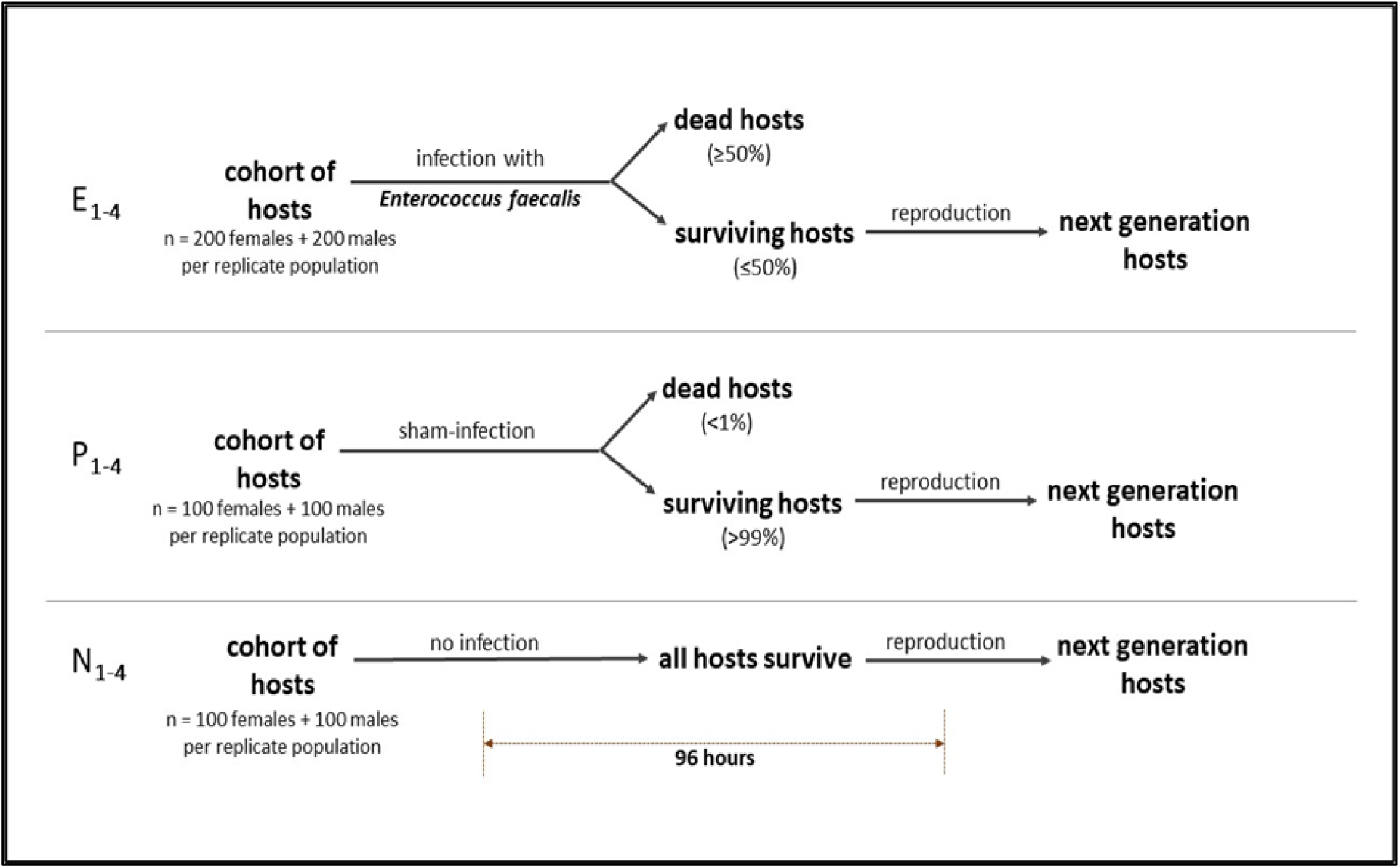
Schematic for the E, P and N selection regimes, each consisting of four replicate populations.

The EPN populations were derived from the ancestral Blue Ridge Baseline (BRB_1-4_) populations. The E_1_, P_1_, and N_1_ populations were derived from BRB_1_, the E_2_, P_2_, and N_2_ populations were derived from BRB_2_, and so on. Selected populations with the same numeric subscript therefore share a recent common ancestor and are part of the same ancestral ‘block’. Blocks are handled on separate days for both regular maintenance and for experiments.

All the populations are maintained on a 16-day discrete generation cycle (day 1 being the day of egg collection). Infections happen on day 12 every generation. Flies are maintained on standard banana-jaggery-yeast medium, and are housed in an incubator at 25 ^O^C with a 12:12 hours LD cycle and 50% relative humidity. For experiments, flies were generated and handled in a fashion that closely resembles the regular maintenance regime for these populations. On day 1, eggs are collected from each of the 12 populations, at a density of 60-80 eggs per vial (25 mm diameter × 90 mm height), in 6-8 ml of standard food medium; 10 vials are set-up per population. The vials are incubated under the constant environmental conditions detailed above, and the eggs develop into adults by day 10. The adults remain in these vials till day 12, when the adult flies are subjected to the corresponding selection regime according to their population identity. By this point of time, all adults are sexually mature and have mated at least once. After being subjected to selection (infected, sham-infected, or uninfected), adult flies are housed in plexiglass cages (14 cm length × 16 cm width × 13 cm height); each population is housed in a separate cage. Each cage is supplied with fresh food medium, poured into a 60 mm Petri plate. The flies remain in these cages for the next 96 hours, with fresh food medium provided to them every 48 hours. On day 16 (96 hours after selection is imposed), each cage is provided with a fresh food plate for the surviving flies to lay eggs on. On day 17 (18 hours after the start of egg laying), eggs are collected from these plates to start the next generation. Day 17 of the previous generation becomes the day 1 of the following generation.

During regular maintenance, there is negligible mortality in the P_1-4_ populations (<1%) and no mortality in the N_1-4_ populations, from the point of handling of adults on day 12 till the start of oviposition window on day 16. During the selection process, the mortality of flies from E_1-4_ populations are maintained at about fifty percent. To this effect, the flies were infected at an infection dose of OD_600_ = 0.8 (see below for more details) between generations 1 and 20 of forward selection. Thereafter, the flies were infected with OD_600_ = 1.0 from generation 21 to 40, with OD_600_ = 1.2 from generation 41 to 60, and with OD_600_ = 1.5 from generation 61 onwards.

### 2.2. Standardization and derivation of experimental flies

Prior to experiments, flies from the three selection regimes were reared for a generation under ancestral maintenance conditions. This is done to account for any non-genetic parental effects (Rose 1984), and flies thus generated are referred to as standardized flies. To generate standardized flies, eggs were collected from all the populations at a density of 60-80 eggs per vial (with 6-8 ml food medium); 10 such vials were set up per population. The vials were incubated under standard maintenance conditions detailed above. On day 12 after egg collection, the adults were transferred to plexiglass cages (14 cm × 16 cm × 13 cm) with food plates (Petri plates, 60 mm diameter). Eggs for experimental flies were collected from these standardized population cages.

### 2.3. Pathogen handling and Infection protocol

*Enterococcus faecalis* (Lazzaro et al., 2006), a Gram-positive bacterium, and a known entomopathogen (Troha and Buchon 2019) was used for infecting the host individuals, both during regular population maintenance and for experiments. All experimental infections were done at an infection dose of OD_600_ = 1.5. A OD_600_ = 1.0 suspension of *E. faecalis* amounts to 10^6^ CFUs per milliliter.

The bacteria are preserved as glycerol stocks (17%) in -80 ^O^C. To obtain live bacterial cells for infections, 10 ml lysogeny broth (Luria Bertani Broth, Miler, HiMedia) is inoculated with glycerol stocks of the bacterium and incubated overnight with aeration (150 rpm in a shaker incubator) at suitable temperature (37 ^O^C). 100 microliters from this primary culture is inoculated into 10 ml fresh lysogeny broth and incubated for the necessary amount of time to obtain confluent (OD_600_ = 1.0-1.2) cultures. The bacterial cells are then pelleted down using centrifugation and resuspended in sterile MgSO_4_ (10 mM) buffer to obtain the required optical density (OD_600_) for infection. Flies are infected, under light CO_2_ anesthesia, by pricking them on the dorsolateral side of their thorax with a 0.1 mm Minutien pin (Fine Scientific Tools, USA) dipped in the bacterial suspension. Sham-infections (injury controls) are carried out in the same fashion, except by dipping the pins in sterile MgSO_4_ (10 mM) buffer.

### 2.4. Systemic pathogen load estimation

In this study, systemic pathogen load was estimated for both dead and alive flies. For estimation of <u>B</u>acterial <u>L</u>oad <u>U</u>pon <u>D</u>eath (BLUD), the dead flies were surface sterilized twice with 70% ethanol (twice for 1 minute) and individually homogenized in 200 microliters of sterile MgSO_4_ (10 mM) buffer within an hour of their death. This homogenate was then serially 1:10 diluted eight times; 10 microliters of each dilution, along with 10 microliters of the original homogenate, was spotted onto Luria Bertani (Miller) agar (2%) plates. The plates were incubated for 8 hours at 37 ^O^C. Colony forming units (CFUs) were counted for the dilution where CFUs ranged between 25 to 250, and the count was multiplied by the appropriate dilution factor to calculate the systemic pathogen load. For estimation of bacterial load of alive flies, a protocol similar to BLUD estimation was followed, except that the living flies were individually homogenized in 50 microliters sterile MgSO_4_ (10 mM) buffer to begin with.

### 2.5. Post-infection survival assay

For assaying post-infection survival, 2–3-day old adult flies from N_1-4_, P_1-4_, and E_1-4_ populations were either infected with *Enterococcus faecalis* (n = 100 males and 100 females per population per block) or sham-infected (n = 50 males and 50 females per population per block), and housed in individual cages (14 cm length × 16 cm width × 13 cm height) with food plates (Petri plates, 60 mm diameter). Fresh food was provided every 48 hours. The survival of these flies were monitored every hour for the first 48 hours and thereafter every 4-6 hours till 96 hours post-infection (HPI). Each block was handled separately on separate days. Survival assay was conducted twice, once after 65 generations of forward selection and again after 75 generations of forward selection.

### 2.6. Female fecundity assay (0–48 hours post-infection)

Fecundity of females during 0–48 HPI was assayed after 70 generations of forward selection. For this assay, 4–5-day old, inseminated females from N_1-4_ and E_1-4_ populations were either infected with *E. faecalis* (n = 80 females per population per block) or sham-infected (n = 40 females per population per block), and thereafter housed individually in food vials where they could oviposit. The survival of these females was monitored every 2 hours till 48 HPI. At the end of this window, all surviving females were discarded, and the vials were incubated under standard maintenance conditions, and the number of progenies eclosing out of these vials were counted 12 days after the end of oviposition period.

### 2.6. Female fecundity assay (96–120 hours post-infection)

Fecundity of females during 96–120 HPI was assayed along with the post-infection survival assay conducted after 75 generations of forward selection. At 96 HPI, alive females from N_1-4_ and E_1-4_ populations, of both infected and sham-infected treatments, were aspirated out of their respective cages and housed individually in food vials (n = 30 females per population per treatment per block). These females were allowed to oviposit for 24 hours, after which they were discarded. The vials with laid eggs were incubated under standard maintenance conditions, and the number of progenies eclosing out of these vials were counted 12 days after the oviposition period. The assay design closely matches the regular maintenance regime, where 96 hours after infection the flies are allowed to oviposit on fresh food medium, and these eggs are used to start the next generation.

### 2.7. Systemic bacterial load of dead flies

Systemic bacterial load of dead flies (BLUD) was assayed along with the post-infection survival assay conducted after 75 generations of forward selection. Dead females and males of N_1-4_ and E_1-4_ populations were aspirated out of the cages within an hour of their death, and their systemic bacterial load was estimated using the above-described protocol. The sample size varied between 15-30 flies per sex per population per block, depending upon how many flies perished due to infection. Since no death was recorded in sham-infected flies, systemic load of sham-infected flies was not monitored during this assay.

### 2.8. Systemic bacterial load of alive flies

To assay for the time dependent changes in systemic bacterial load in living, infected females, 700 N_1-4_ females and 350 E_1-4_ females (per block) were infected with *E. faecalis*, and housed in separate cages. Starting sample size of N females was double of that of the E females to ensure enough surviving flies were available for bacterial load measurements at later time points, given that N females are expected to have much greater mortality compared to E females. Starting from 3 hours post-infection (HPI), every 3 hours, the systemic bacterial load of the females were assayed for the first 48 hours, after which the sampling frequency was reduced to 6-12 hours. The assay continued till 96 HPI. At every sampling time point, 10 alive females of each population were aspirated out of their respective cages, and their systemic bacterial load was estimated using the above-described protocol. Individual blocks were assayed on separate days. To assay for the time dependent changes in systemic bacterial load in living, infected males, we followed an identical experimental design, except that the starting sample sizes for N_1-4_ and E_1-4_ males were 240 and 120 (per block), respectively, and measurement of systemic bacterial load was carried out only at 4-, 10-, 48-, and 96-HPI. This assay was done for females after 78 generations of forward selection and for males after 80 generations of forward selection. Systemic bacterial load for sham-infected flies were measured only at one time point: 3 HPI for females and 4 HPI for males. No sham-infected fly yielded any CFU during any of the experiments.

### 2.9. Statistical analysis

All statistical analyses were carried out using R statistical software (version 4.1.0, R Core Team 2021). Post-infection survival of infected flies was analyzed using a mixed-effects Cox proportional hazards model, which included host selection history, host sex, and their interaction as fixed factors, and block as a random factor. The model was subjected to Analysis of Deviance (type II) for significance testing. Analysis was done for the survival assays carried out in generations 65 and 75 separately, and only the infected flies were considered since there was negligible mortality in the sham-infected flies. Female fecundity was analyzed using Analysis of Variance (type III ANOVA), with infection treatment, host selection history, and their interaction as fixed factors, and block as a random factor. Bacterial Load Upon Death (BLUD, log2 transformed systemic bacterial load) was analyzed using ANOVA (type III), with time of death (HPI), host evolutionary history, host sex, and their interaction as fixed factors, and block and its interaction with the fixed factors as random factors. Post-hoc pairwise comparisons for ANOVA were carried out using Tukey’s HSD method. Bacterial load in living flies (log2 transformed systemic bacterial load) was analyzed using ANOVA (type III), with hours post-infection (HPI), host evolutionary history, and their interaction as fixed factors, and block and its interaction with the fixed factors as random factors. Data for each sex was analyzed separately. Pairwise comparison between bacterial load carried by E and N flies at each HPI was carried out using Holm-Sidak method for p-value correction.

## 3. Results

### 3.1. Post-infection survival

To test for the response to selection, flies from the N_1-4_, P_1-4_, and E_1-4_ populations were infected with *Enterococcus faecalis*, and their post-infection survival was measured. Post-infection survival assay was conducted twice: once after 65 generations of forward selection and again after 75 generations of forward selection.

When assayed for after 65 generations of forward selection, selection history had a significant effect on post-infection survival (**table 1.A**). Host sex, and selection history × host sex interaction did not have any effect on post-infection survival (**table 1.A**). E females (hazard ratio, 95% confidence interval: 0.330, 0.263-0.415) exhibited better post-infection survival compared to N females; P females (HR, 95% CI: 0.949, 0.791-1.139) did not differ from N females in terms of post-infection survival (**figure 2.A**). Similarly, E males (HR, 95% CI: 0.305, 0.241-0.386) exhibited better post-infection survival compared to N males; P males (HR, 95% CI: 0.943, 0.785-1.134) did not differ from N males in terms of post-infection survival (**figure 2.B**).

**Figure 2.**
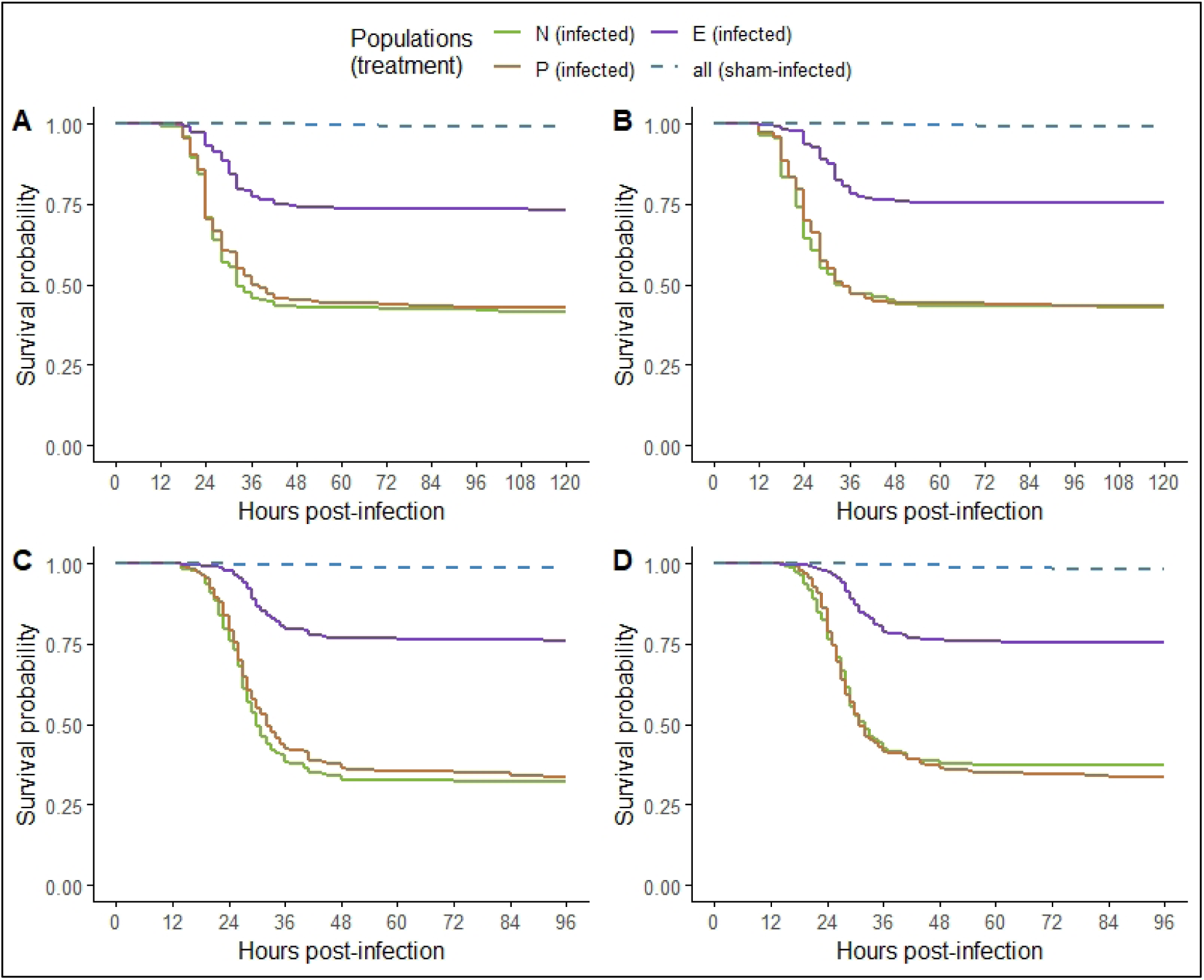
Survival of flies from the N_1-4_, P_1-4_, and E_1-4_ populations after being infected with *Enterococcus faecalis* or being sham-infected. Survival curves plotted using Kaplan-Meier method after pooling data across all four replicates for each selection regime. **(A)** Survival of females, generation 65. **(B)** Survival of males, generation 65. **(C)** Survival of females, generation 75. **(D)** Survival of males, generation 75.

**Table 1.**
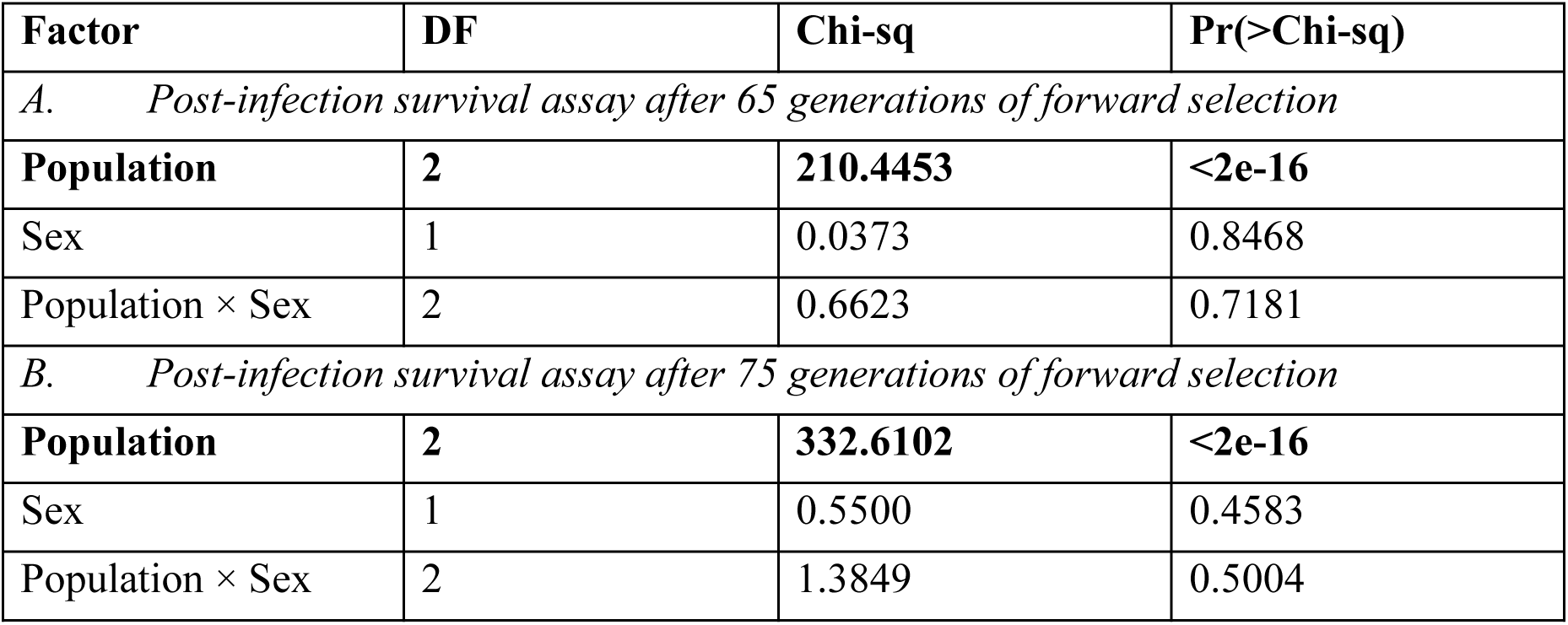
Analysis of deviance (type II) on mixed-effect Cox proportional hazards model to test for the effect of selection history (population identity), sex, and their interaction on survival of flies after being infected with *Enterococcus faecalis*. Significant effects (p < 0.05) are marked in bold font.

When assayed for after 75 generations of forward selection, selection history had a significant effect on post-infection survival (**table 1.B**). Host sex, and selection history x host sex interaction did not have any effect on post-infection survival (**table 1.B**). E females (HR, 95% CI: 0.220, 0.174-0.278) exhibited better post-infection survival compared to N females; P females (HR, 95% CI: 0.908, 0.767-1.076) did not differ from N females in terms of post-infection survival (**figure 2.C**). Similarly, E males (HR, 95% CI: 0.250, 0.198-0.317) exhibited better post-infection survival compared to N males; P males (HR, 95% CI: 1.026, 0.862-1.221) did not differ from N males in terms of post-infection survival (**figure 2.D**).

### 3.2. Systemic bacterial load in dead flies

<u>B</u>acterial <u>L</u>oad <u>U</u>pon <u>D</u>eath (BLUD) of infected females and males from N_1-4_ and E_1-4_ populations was measured along with the post-infection survival assay conducted after 75 generations of forward selection. Only host sex had a significant effect on BLUD (**table 2.C**); time of death, selection history, and selection history × host sex did not have a significant effect on BLUD (**figure 3**). Pair-wise comparison using Tukey’s HSD (**supplementary table S1.C**) indicated that N males (Least-square mean, 95% CI: 24.5, 23.8-25.2) and E males (LS mean, 95% CI: 24.2, 23.4-25.0) carried similar bacterial loads at the time of their death, and so did N females (LS mean, 95% CI: 25.9, 25.2-26.6) and E females (LS mean, 95% CI: 25.8, 25.1-26.6) females. Pooled across both populations, the males carried significantly lower bacterial loads at the time of their death compared to the females.

**Figure 3.**
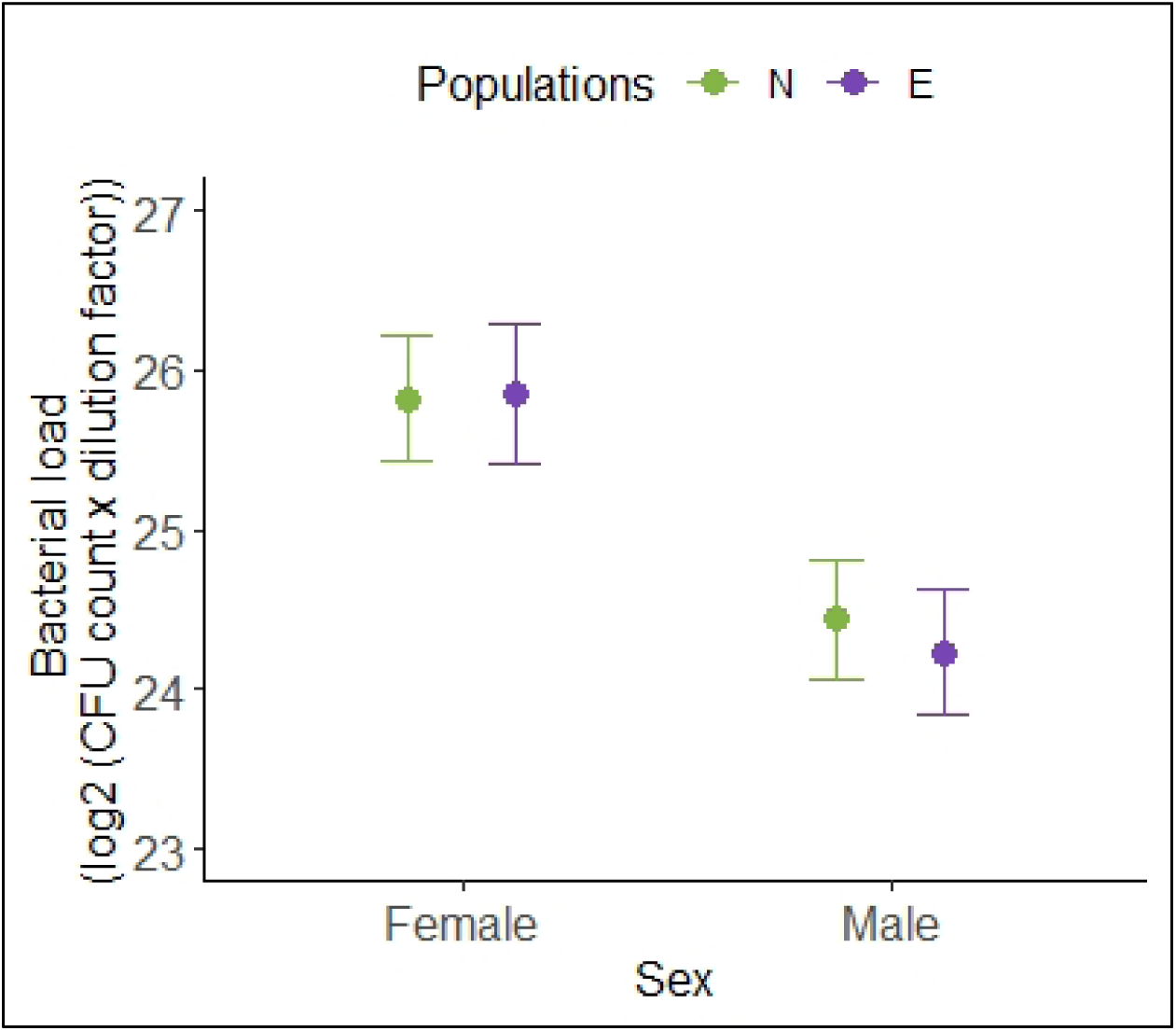
<u>B</u>acterial <u>L</u>oad <u>U</u>pon <u>D</u>eath (BLUD) of flies from the N_1-4_ and E_1-4_ populations infected with *Enterococcus faecalis*. Data pooled across all four replicates for each selection regime. Y-axis error bars represent 95% confidence intervals around the respective means.

**Table 2.** Analysis of variance (ANOVA, type III) for the effect of selection history (population identity), sex, etc. on various assayed traits. Significant effects (p < 0.05) are marked in bold font. (HPI: hours post-infection.)

| Factors | SS | MS | DF | Residual DF | F value | p-value |
| --- | --- | --- | --- | --- | --- | --- |
| *A. Female fecundity, during 0—48 hours post-infection* |  |  |  |  |  |  |
| Treatment | 0.01506 | 0.01506 | 1 | 4.04 | 0.0826 | 0.78790 |
| Population | 0.28966 | 0.28966 | 1 | 3.99 | 1.5891 | 0.27607 |
| **Treatment × Population** | **0.78827** | **0.78827** | **1** | **946.36** | **4.3246** | **0.03783** |
| *B. Female fecundity, during 96—120 hours post-infection* |  |  |  |  |  |  |
| **Treatment** | **1697.95** | **1697.95** | **1** | **462** | **15.976** | **7.467e-05** |
| **Population** | **720.51** | **720.51** | **1** | **462** | **6.779** | **0.00952** |
| Treatment × Population | 188.66 | 188.66 | 1 | 462 | 1.775 | 0.18342 |
| *C. Bacterial load upon death (BLUD)* |  |  |  |  |  |  |
| HPI | 0.494 | 0.494 | 1 | 296 | 0.1311 | 0.717528 |
| Population | 1.037 | 1.037 | 1 | 6 | 0.2752 | 0.619233 |
| **Sex** | **92.595** | **92.595** | **1** | **4** | **24.5787** | **0.007409** |
| Population × Sex | 1.013 | 1.013 | 1 | 4 | 0.2690 | 0.632114 |
| *D. Within-host bacterial dynamics, females* |  |  |  |  |  |  |
| **HPI** | **527.88** | **527.88** | **1** | **84** | **35.611** | **5.549e-08** |
| **Population** | **1842.30** | **1842.30** | **1** | **20** | **124.281** | **5.495e-10** |
| **HPI × Population** | **562.32** | **562.32** | **1** | **1661** | **37.914** | **9.159e-10** |
| *E. Within-host bacterial dynamics, males* |  |  |  |  |  |  |
| **HPI** | **109.52** | **109.52** | **1** | **320** | **23.091** | **2.38e-06** |
| **Population** | **373.66** | **373.66** | **1** | **320** | **78.785** | **< 2.2e-16** |
| **HPI × Population** | **107.72** | **107.72** | **1** | **320** | **22.712** | **2.86e-06** |

### 3.3. Systemic bacterial load in living flies

Systemic bacterial load of living, infected females from N_1-4_ and E_1-4_ populations was measured after 78 generations of forward selection. Time of sampling (hours post-infection), selection history, and the interaction between the two had a significant effect on the bacterial load carried by living females (**table 2.D**), with E females having either less or equal bacterial load compared to N females at every sampling point (**figure 4.A**). Using Holm-Sidak correction of p-values (**supplementary table S2.A**), we identified that E females carried significantly less bacterial load compared to N females at 3- to 24-, 30-, and 42-hours post-infection (HPI).

**Figure 4.**
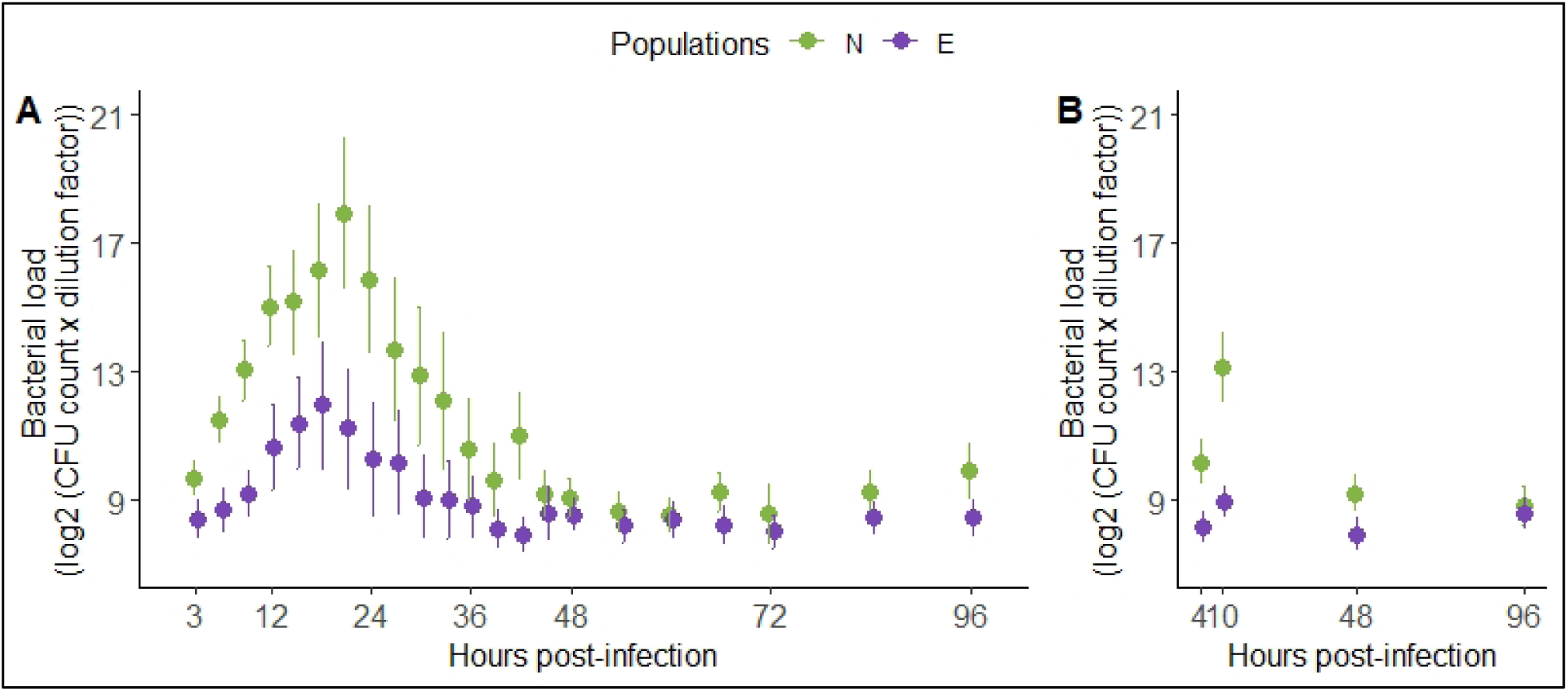
Systemic bacterial load carried by alive flies from the N_1-4_ and E_1-4_ populations at different time-points (hours post-infection) after being infected with *Enterococcus faecalis*. Data pooled across all four replicates for each selection regime. Y-axis error bars represent 95% confidence intervals around the respective means. **(A)** Systemic bacterial load of females. **(B)** Systemic bacterial load of males.

Systemic bacterial load of living, infected males from N_1-4_ and E_1-4_ populations was measured after 80 generations of forward selection. Time of sampling (hours post-infection), selection history, and the interaction between the two had a significant effect on the bacterial load carried by living males (**table 2.E**), with E males having either less or equal bacterial load compared to N males at every sampling point (**figure 4.B**). Using Holm-Sidak correction of p-values (**supplementary table S2.B**), we identified that E males carried significantly less bacterial load compared to N males at 4-, 10-, and 48-hours post-infection (HPI).

### 3.4. Female fecundity (0-48 hours post-infection)

Female fecundity (the number of progeny produced per hour by an individual female) during 0-48 hours post-infection of infected and sham-infected females from N_1-4_ and E_1-4_ populations was measured after 70 generations of forward selection. Infected females succumbed to infection during the period in which fecundity was being measured in this assay. Hence, to accommodate differences in survival times between infected and sham-infected females, and amongst infected females, the number of progeny produced by an individual female was divided by the number of hours the female survived, and this standardized number of progeny produced per hour was used as the measure of fecundity in this assay.

During the assay period, no mortality was recorded in the sham-infected females from either N_1-4_ or E_1-4_ populations (**figure 5.A**). Among the infected females, females from the E_1-4_ populations died significantly less compared to females from the N_1-4_ populations (HR, 95% CI: 0.292, 0.231-0.369). Neither selection history nor infection treatment had any significant effect on female fecundity, but selection history × infection treatment had a significant effect (**table 2.A**, **figure 5.B**). Pairwise comparison using Tukey’s HSD (**supplementary table S1.A**) indicated that E-infected females (least-square mean, 95% CI: 0.587, 0.308-0.867) produced significantly less number of progeny per hour compared to N-infected (LS mean, 95% CI: 0.686, 0.407-0.966), E-sham-infected (LS mean, 95% CI: 0.733, 0.455-1.011), and N-sham-infected (LS mean, 95% CI: 0.712, 0.434-0.990) females. N-infected, E-sham-infected, and N-sham-infected females did not differ from one another in terms of the number of progeny produced per hour.

**Figure 5.**
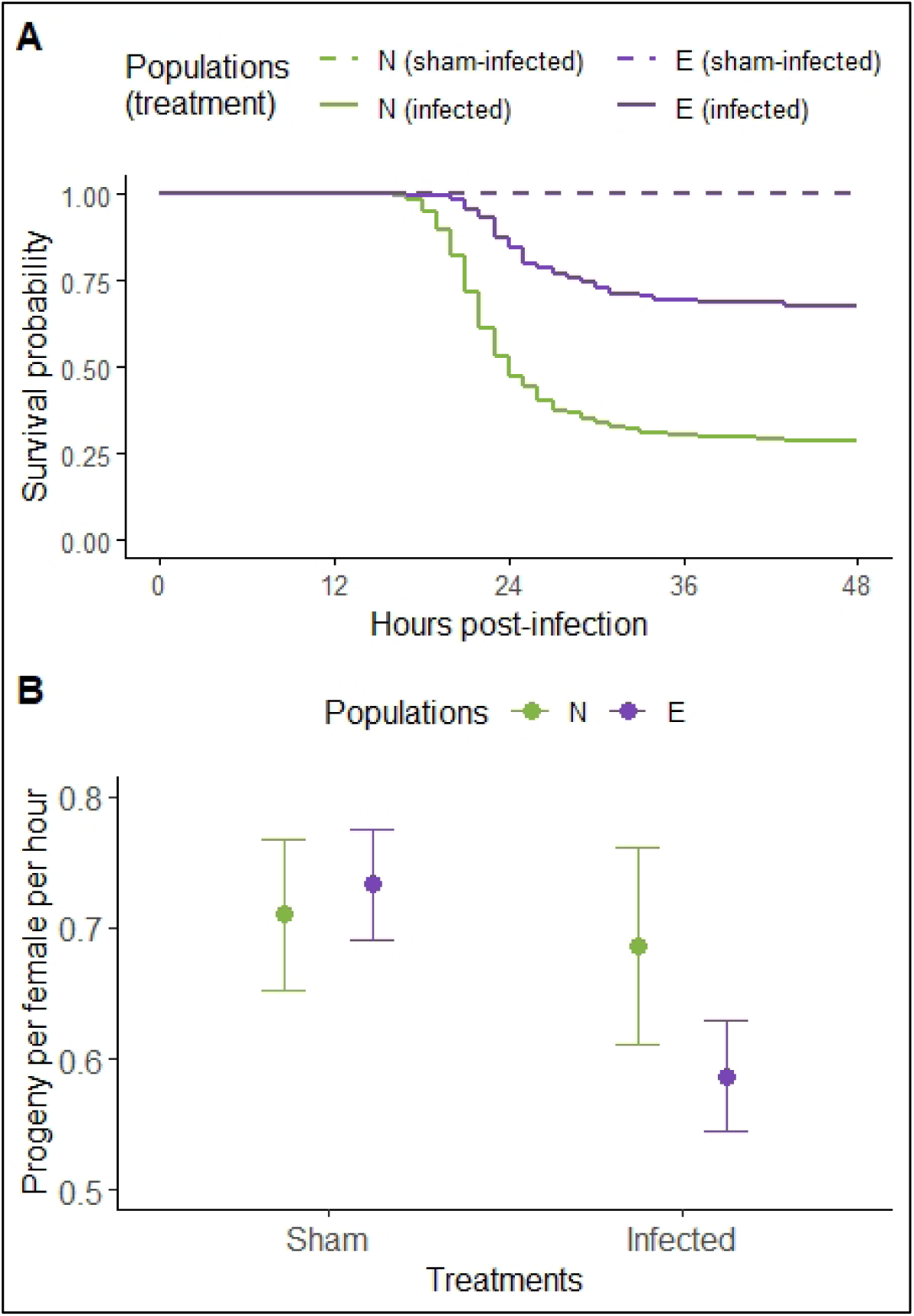
Survival and fecundity of females from the N_1-4_ and E_1-4_ populations during 0–48 hours post-infection. **(A)** Survival of the females after being subjected to infection with *Enterococcus faecalis* or to sham-infection. Survival curves plotted using Kaplan-Meier method after pooling data across all four replicates for each selection regime. **(B)** Fecundity of females after being subjected to infection with *Enterococcus faecalis* or to sham-infection. Data pooled across all four replicates for each selection regime. Y-axis error bars represent 95% confidence intervals around the respective means.

### 3.5. Female fecundity (96-120 hours post-infection)

Female fecundity (the number of progeny produced by an individual female) during 96-120 hours post-infection of infected and sham-infected females from N_1-4_ and E_1-4_ populations was measured along with the post-infection survival assay conducted after 75 generations of forward selection. Since no death was recorded during the assay period in females of either infection treatment, absolute number of progeny produced by an individual female was used as measure of fecundity in this assay.

Selection history and infection treatment had a significant effect on female fecundity (**table 2.B**), but their interaction did not have a significant effect (**figure 6**). Pairwise comparison using Tukey’s HSD (**supplementary table S1.B**) indicated that N-infected females (LS mean, 95% CI: 22.7, 19.4-26.1) produced a significantly greater number of progeny compared to N-sham-infected (LS mean, 95% CI: 17.6, 14.3-20.9), E-infected (LS mean, 95% CI: 18.9, 15.6-22.2), and E-sham-infected (LS mean, 95% CI: 16.4, 13.1-19.7) females. N-sham-infected, E-infected, and E-sham-infected females did not differ from one another in terms of the number of progeny produced.

**Figure 6.**
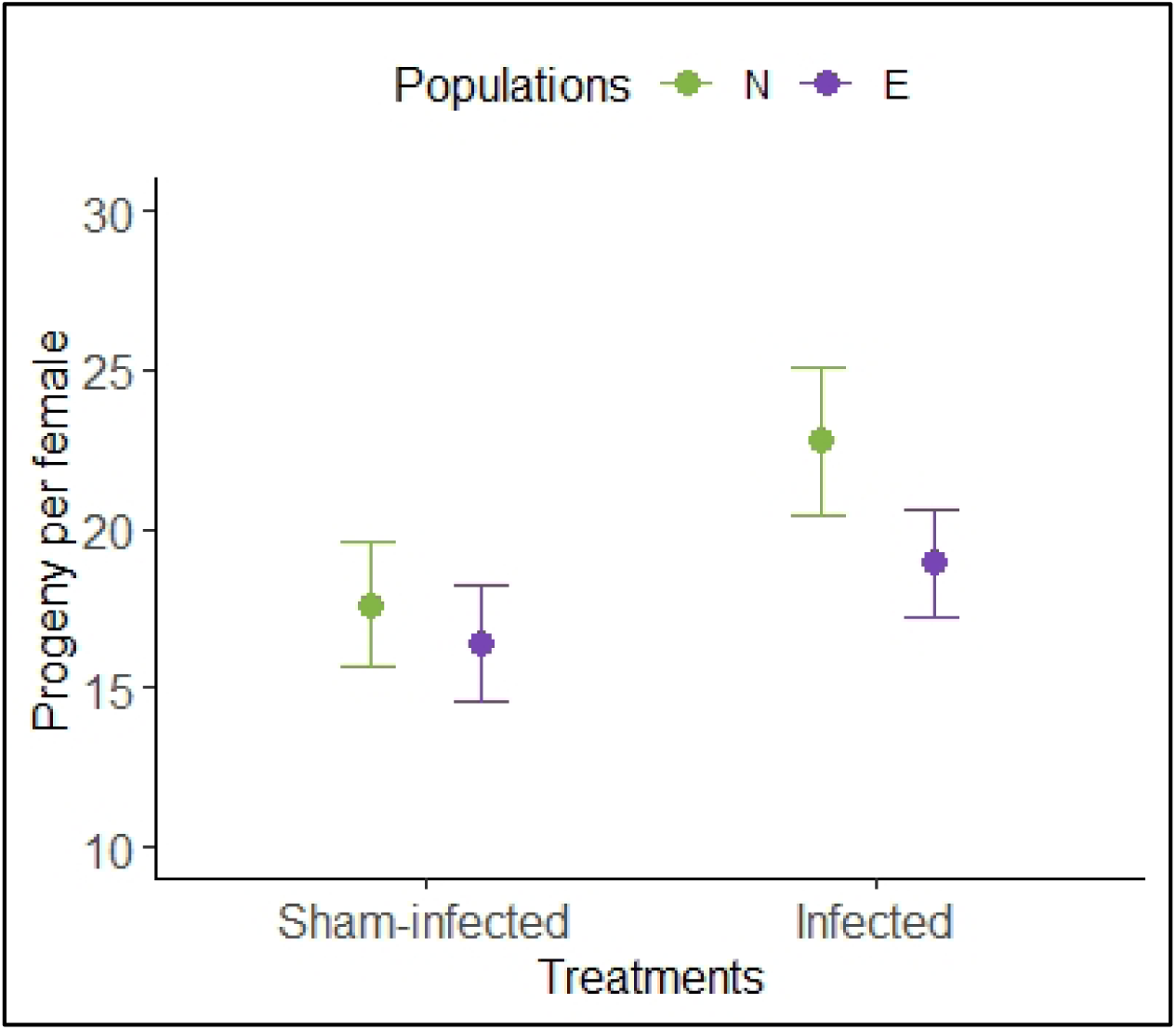
Fecundity of females from the N_1-4_ and E_1-4_ populations during 96–120 hours post-infection, after being subjected to infection with *Enterococcus faecalis* or to sham-infection. Data pooled across all four replicates for each selection regime. Y-axis error bars represent 95% confidence intervals around the respective means.

## 4. Discussion

The fitness of an infected host is contingent upon both its resistance and tolerance to the infecting pathogen. Host resistance determines the intensity of the infection (i.e., the systemic pathogen load carried by the host), while host tolerance determines the effect of a given infection intensity on host fitness (i.e., the level of morbidity or mortality experienced by the host). In this study, we investigated whether hosts evolve increased resistance, tolerance, or both when subjected to selection for increased post-infection survival in a laboratory experimental evolution set-up in which host reproduction is contingent upon the host successfully surviving the infection.

Briefly, we experimentally evolved replicate *Drosophila melanogaster* populations for increased post-infection survival after being infected with an entomopathogenic Gram-positive bacterium, *Enterococcus faecalis*. The strain of *E. faecalis* used for imposing selection pressure on the hosts was originally isolated from wild-caught flies (Lazzaro et al., 2006), making it suitable for use in a study of experimental evolution of immune function and defense strategies. The selected flies were infected with the bacterium every generation, and only those individuals that survived the infection were allowed to reproduce and contribute to the next generation (**figure 1**). Within 35 generations of forward selection, the flies of the selected (E_1-4_) populations exhibited increased post-infection survival compared to the flies of both the procedural control (P_1-4_) and the uninfected control (N_1-4_) populations (Singh et al., 2021). The selected populations continued to become significantly better at surviving infection with *E. faecalis* with continued forward selection, as demonstrated by the results from the post-infection survival assays reported here, carried out after 65 generations (**figures 2.A and 2.B**) and 75 generations (**figures 2.C and 2.D**) of forward selection. The procedural control and uninfected control populations exhibited similar post-infection survival in all assays across generations (**figure 2**). Given that the selected populations rapidly evolved increased post-infection survival, we explored if this increase in survival in these populations was explained by increase in disease resistance, disease tolerance, or both.

### 4.1. No evidence for evolution of tolerance

To compare the tolerance of flies from the selected and control populations, we compared the <u>B</u>acterial <u>L</u>oad <u>U</u>pon <u>D</u>eath (BLUD) of infected females and males from the E_1-4_ and N_1-4_ populations. BLUD has been suggested as a suitable proxy of host tolerance to bacterial infections in *D. melanogaster* in previous studies (Duneau et al., 2017a, Duneau et al., 2017b, Vincent et al., 2020). The results show that the flies from the selected (E_1-4_) and the control (N_1-4_) populations carry similar BLUD (**figure 3**), suggesting that the flies succumb to infection at similar systemic pathogen loads irrespective of their selection history. We propose that this observation indicates that there has been no evolution of host tolerance in the selected populations, even after 75 generations of forward selection.

The results also show that there is sexual dimorphism in BLUD: females of both the selected and the control populations carry a significantly higher BLUD compared to males of their respective populations (**figure 3**). This observation may represent sexual dimorphism in tolerance to infection with *E. faecalis* in these populations, with the females being more tolerant to *E. faecalis* infection than the males. Sexual dimorphism in disease tolerance is known in *D. melanogaster* (Vincent and Sharp 2014), although a previous study has reported that BLUD for *Providencia rettgeri* is not affected by host sex in certain laboratory lines of this species (Duneau et al., 2017a). Alternatively, our observation may simply be a reflection of sexual dimorphism in body size. *D. melanogaster* females tend to be bigger compared to males, and this trend holds true in our populations too, as has been demonstrated by previous experiments (Singh et al., 2022). It is therefore possible that the females, by virtue of having a greater body size compared to the males, succumb to infection, and die at a greater systemic bacterial load, one must note that the difference in body size previously reported is much less in relative magnitude than the difference in BLUD observed here.

### 4.2. Evolution of increased resistance

A more resistant host, by definition, is better at limiting systemic proliferation of pathogens. Therefore, we measured the systemic bacterial load of living flies from the E_1-4_ and N_1-4_ populations at regular intervals after being infected with *E. faecalis*, to study the dynamics of systemic bacterial growth during different stages of infection. The results demonstrate that in the infected females (**figure 4.A**) of both E_1-4_ and N_1-4_ populations, systemic bacterial load initially increases (3—18 hours post-infection, HPI) and then decreases (18—48 HPI), and thereafter eventually settles down to a somewhat stagnant bacterial load (48—96 HPI). Mortality due to *E. faecalis* infection in these populations is mostly limited to before 48 HPI, with very few flies dying after 48 HPI (**figure 2**). Therefore, 0—48 HPI represents the *acute phase* of *E. faecalis* infection in these populations, and the period after 48 HPI represents the *chronic phase* (*sensu* Chambers et al., 2019, Hidalgo et al., 2022).

Females of the selected (E_1-4_) populations carry a lower systemic bacterial load compared to females of the control (N_1-4_) populations during the acute phase of infection (**figure 4.A**); significantly so during 3—30 HPI (**supplementary table S2.A**). From 48 HPI onwards, females from both populations carry a similar <u>S</u>et <u>P</u>oint <u>B</u>acterial <u>L</u>oad (SPBL, *sensu* Duneau et al., 2017b) during the chronic phase of infection. Similar dynamics is seen in case of systemic bacterial load in males, with males of the selected (E_1-4_) populations carrying a lower systemic bacterial load compared to males of the control (N_1-4_) populations during 4—48 HPI, but having similar SPBL at 96 HPI (**figure 4.B, supplementary table 2.B**). These observations suggest that the E_1-4_ flies are better at controlling the proliferation of bacteria within their body compared to N_1-4_ flies. We therefore propose that the selected populations have evolved increased resistance to *E. faecalis* infection in response to the selection for increased post-infection survival imposed upon them for 78—80 generations.

Our results further indicate that the difference between systemic bacterial loads in infected flies from E_1-4_ and N_1-4_ populations is evident at the earliest time point for which bacterial load was measured in our experiments (3 HPI for females and 4 HPI for males). Previous studies have proposed that systemic bacterial dynamics early on in infection eventually determine infection outcome at later stages (Duneau et al., 2017b). An early difference in the systemic bacterial loads carried by the flies from the selected and the control populations might additionally suggest that whatever mechanism underlies the increased resistance in the selected populations is either a constitutively active defense or a defense that can be rapidly induced following infection. Further studies are required to test out this hypothesis.

### 4.3. Fecundity cost associated with increased resistance

Previous studies, both theoretical and empirical, have suggested that evolution of increased resistance imposes various costs on the host organism (reviewed in Raberg and Stjernman 2012). Such costs may manifest as physiological trade-offs, such as reduced reproductive capacity, both in presence and in absence of an infection (Schmid-Hempel 2005, McKean and Lazzaro 2011). Therefore, we investigated if evolution of increased resistance in the selected populations is accompanied by a concomitant reduction in female fecundity, with and without being infected with *E. faecalis*. We compared fecundity of infected and sham-infected females from the E_1-4_ and N_1-4_ populations, both during the acute phase (0—48 HPI) and chronic phase (96—120 HPI) of infection, in two separate experiments. The effect of bacterial infections on female fecundity is known to change depending upon the infection phase in *D. melanogaster* (Howick and Lazzaro 2014). Therefore, examining the presence of costs during both the acute and the chronic phase is important. It should be noted that the window between 96—120 HPI also coincides with the breeding window of these populations during their regular maintenance (**figure 1**), and therefore is to be considered as their fitness window. During regular population maintenance, only the flies that reproduce during this period contribute to future generations, and therefore, fecundity of females during this period is akin to their lifetime reproductive success.

When female fecundity was measured during the acute phase of infection (**figure 5.B**), females from the selected (E_1-4_) and the control (N_1-4_) populations had similar fecundity when sham-infected, i.e., in absence of infection. This observation suggests that the selected populations do not pay any *maintenance cost* (*sensu* McKean et al 2008, McKean and Lazzaro 2011) of increased resistance. When infected with *E. faecalis*, females from the selected populations exhibited a decline in fecundity compared to sham-infected females. Such a decline in fecundity was not observed in case of females from the control populations. These observations suggest that the selected populations incur an *evolved cost of immune deployment* (*sensu* McKean et al 2008, McKean and Lazzaro 2011) associated with their increased resistance. This decline in fecundity of females from the selected populations after being infected can be caused either by resource/energy trade-offs between reproduction and defense mechanisms (Schmid-Hempel 2003), or by damage to the reproductive tissue from the infection (Brandt and Schneider 2006).

When female fecundity was measured during the chronic phase of infection (**figure 6**), the females from the selected (E_1-4_) populations had similar fecundity irrespective of whether they were infected with *E. faecalis* or sham infected. The females from the control (N_1-4_) populations exhibited a mild, but significant, increase in fecundity after being infected. Again, in absence of infection, the females from the selected and the control populations had similar fecundity. Therefore, during the chronic phase of infection, we do not see any cost of increased resistance, maintenance or deployment, in the selected populations. Importantly, since this window of fecundity measurement also represents the selection window of these populations, during the *selection window*, when it matters most, the selected populations incur no fecundity cost of increased resistance. Additionally, absence of a cost of immune maintenance (i.e., no cost of improved resistance in absence of infection) during both the acute and chronic phase of infection makes it unlikely that the selected hosts in our study have evolved a constitutively active defense mechanism.

Absence of any fecundity costs during the selection window has two important implications. *One*, this shows that flies can recover from suppression of fecundity induced by *E. faecalis* infection and that chronic infection with this pathogen does not exact a fitness cost on the host. This also suggests that the suppression of fecundity observed during the acute phase is more likely to be driven by resource reallocation trade-off and not damage to the reproductive tissue, assuming that such damages are permanent in nature. And *two*, absence of costs associated with increased resistance facilitates the evolution of resistance. Physiological costs associated with disease resistance is one of the major factors that limit evolution of increased resistance under natural settings (Raberg & Stjernman 2012). We propose that this apparent absence of physiological costs of resistance to *E. faecalis* during the selection window therefore might have favored the evolution of increased resistance in our selected populations, and may have also relaxed the selection for evolution of tolerance.

Additionally, presence of a cost of deployment in the selected populations only during the acute phase of infection has an interesting implication. In principle, it is possible that the females from the selected populations have evolved fecundity suppression as a defense strategy: they temporally partition their priorities. Early in infection, during the acute phase, the selected females reduce reproductive investment and re-allocate available resources towards resisting systemic pathogen growth, thereby improving resistance. Following this period, they recover from the infection, and thereafter in the selection window (which coincides with the chronic phase of infection), they invest towards reproduction and thereby ensuring their contribution to the next generation. Previous studies have debated if post-infection suppression of fecundity is a consequence of the pathogen manipulating the host physiology or a host defense strategy to better its own fitness (Hurd 2001). It has also been previously questioned if suppression of reproductive effort can indeed help hosts improve their post-infection survival (Javois 2013). Therefore, we propose that our observed results acts as a proof-of-principle that suppression of fecundity can indeed help the host survive an infection challenge, and can also evolve as a viable defense strategy.

In addition to absence of costs associated with increased resistance, evolution of tolerance in the selected populations may have been hindered by lack of heritable genetic variation for tolerance. Previous studies have argued that wild populations are not likely to harbor genetic polymorphism for tolerance, because whenever a more tolerant mutant invades a population, it rapidly goes to fixation (Roy and Kirchner 2000). Therefore, although the baseline populations used to initiate the selected populations in our study were derived from wild-caught flies (Singh et al., 2015), it is possible that there was a lack of heritable genetic variation for tolerance to *E. faecalis* infection in the starting populations. However, we think that this is an unlikely scenario: these same baseline populations have been successfully used to initiate three separate experimental evolution set-ups, all selecting for increased post-infection survival, including the present populations (Gupta et al., 2016, Ahlawat et al., 2022, Singh et al., 2021). At every instance, a rapid response to selection was observed, suggesting an abundance of heritable genetic variation for antibacterial defense strategies. However, since partitioning of resistance and tolerance was never carried out in the case of these other populations (Gupta et al., 2015, Ahlawat et al., 2022), we cannot confirm if the observed increase in post-infection survival in those populations was due to increased resistance or increased tolerance.

### 4.4. Summary, hypotheses, and speculations

Therefore, to summarize, we have demonstrated in this study that hosts evolve increased disease resistance to bacterial infections when they are subjected to selection for increased post-infection survival following a systemic pathogenic infection with *Enterococcus faecalis*. The same selection pressure does not lead to an evolution of increased disease tolerance. This is a surprising result given that numerous theoretical studies exist that suggest that tolerance as a defense strategy has multiple associated advantages over resistance, and therefore, is more likely to evolve in response to selection imposed by pathogenic infections (reviewed in Raberg & Stjernman 2012). There can be three obvious explanations for the result obtained in this study. *One*, the result is unique to the pathogen strain and the host population used for the experimental evolution set-up, and therefore, is not representative of other experimental evolution set-ups or of any natural setting. *Two*, as discussed above, the ancestral populations used here for experimental evolution lacked heritable variation for tolerance. And *three*, evolution of increased resistance had no associated costs for this particular pathogen in these populations in this experimental evolution set-up. We did find some empirical evidence supporting the third conjecture, although the other two possibilities cannot be completely discounted.

Beyond these three plausible explanations for no evolution of host tolerance in our study, we speculate that the experimental evolution set-up we implemented might have had certain idiosyncrasies that could potentially bias the evolutionary trajectory of the host populations towards preferential evolution of resistance. Below we discuss these idiosyncrasies, which we propose are not unique to our study design, but can be generalized to other similar experimental evolution studies.

#### Separation of mortality and selection windows

Under natural settings the fitness of an organism is determined by its contribution to the next generation, measured as the lifetime reproductive success (Fisher 1930). An infected host can thus maximize its fitness by either surviving the infection, or increasing its immediate reproductive effort, or if it is physiologically feasible, then both. And therefore, under natural settings, strategies like terminal investment (Minchella 1985) and fecundity compensation (Parker et al 2011) can evolve in response to selection from pathogen pressure. In some sense, both these strategies help hosts maximize their reproductive fitness, without affecting host resistance to pathogens directly, and thus should be considered as part of the host tolerance repertoire (Vale and Little 2012, Kutzer and Armitage 2016, Kutzer et al., 2019). Contrary to the natural environment, during laboratory experimental evolution, host reproduction is often made conditional on the host surviving the infection. By design in most, if not all, experimental evolution studies the reproduction window comes after the first wave of infection-induced mortality (the acute phase of infection) has passed (viz. Ye et al., 2009, Martins et al., 2013, Faria et al., 2015, Gupta et al., 2015, Ahlawat et al., 2022). The same is true for our populations. Under such a scenario resistance becomes the only feasible strategy. Evolving to limit systemic pathogen load, and thereby minimizing the somatic damage caused by pathogen-derived virulence factors, helps the infected host survive the infection and reproduce adequately during the selection window. In fact, having the reproduction window after the mortality window can even lead to hosts temporally sorting out their priorities - first resist, then reproduce - as suggested by our results (**figures 5.B and 6**).

#### Maintenance of pathogen prevalence via artificial infection

Under natural settings pathogen presence fluctuates with fluctuations in levels of host resistance. As resistance evolves in the host population, pathogen prevalence falls, thereby also reducing the selection pressure on the host to become more resistant (Roy and Kirchner 2000). Under such circumstances, the cost-to-benefit ratio for increasing resistance becomes steep, and hosts evolve to be less resistant. This again increases pathogen prevalence, which brings back the selection pressure on the hosts to become more resistant. This cyclic process maintains polymorphism in host resistance traits, and resultantly resistance strategies never go to fixation in the host population (Roy and Kirchner 2000). But during laboratory experimental evolution, hosts are artificially infected every generation, thereby maintaining pathogen prevalence. This severs the link between host resistance and pathogen prevalence, thereby ensuring that selection for increased resistance is always maintained irrespective of how resistant the host becomes. Under such a scenario, continuously increasing resistance may be the only feasible strategy available to the host, till the point when resistance cannot be increased further because of physiological constraints. We hypothesize that hosts would evolve tolerance in this type of an experimental evolution set-up only after a prolonged period of selection: when resistance has evolved to its physiological limit or when resistance has evolved to the point that cost of immunopathology outweighs the benefits of pathogen control (Lazzaro and Tate 2022).

#### Absence of antagonistic coevolution and associated increase in pathogen virulence

A major cost associated with evolution of increased resistance under natural settings is that increased resistance in the hosts selects for more virulent pathogens (Miller et al., 2006). But, barring some exceptions (viz. Berenos et al., 2011, Rafaluk-Mohr et al., 2018, Biswas et al., 2018, Ahlawat et al., 2022), in most laboratory experimental studies, especially those using insects as hosts, the pathogen is not allowed to co-evolve with the host (viz. Ye et al., 2009, Martins et al., 2013, Faria et al., 2015, Gupta et al., 2015), including our populations. This ensures that the pathogen virulence remains constant irrespective of however resistant the host becomes, thereby eliminating one of the key costs associated with evolution of increased resistance. Under such a scenario, the host is free to evolve increased resistance *ad libitum*, that is until the physiological capacity to do so is reached. The studies that have explored host-pathogen coevolution using laboratory insect populations, to our knowledge, did not specifically test for evolution of tolerance in the evolved populations (Berenos et al., 2011, Rafaluk-Mohr et al., 2018, Biswas et al., 2018, Ahlawat et al., 2022).

We therefore hypothesize that the evolution of disease tolerance would be observable in an experimental evolution setting where the pathogen prevalence is allowed to fluctuate, and pathogen virulence is allowed to counter evolve as the host becomes better defended against the pathogen. Additionally, the study design should allow for manifestation of the physiological costs associated with evolution of different defense strategies. Since all three of these features were absent from our experimental evolution set-up, we speculate that our selected populations might have been biased towards evolving disease resistance as the preferred strategy of defense.

## Author contributions

AB conceptualized and designed the study; AB, TK, AS, and PND executed the experiments and acquired the data; AB analyzed and interpreted the data; AB and NGP drafted the manuscript; NGP acquired funding for the project. All authors have read, reviewed, and approved the final version of the manuscript.

## Acknowledgement

The authors thank Prof. Brian Lazzaro (Cornell University, USA) for providing the *Enterococcus faecalis* isolate used in the experiments.

## Supplementary tables

**Supplementary table 1.** Pairwise comparisons using Tukey’s HSD accompanying type III ANOVA for the effect of selection history (population identity), sex, etc. on various assayed traits. Significant effects (p < 0.05) are marked in bold font.

| Comparison | Estimate | SE | DF | t ratio | p-value |
| --- | --- | --- | --- | --- | --- |
| *A. Female fecundity, during 0—48 hours post-infection* |  |  |  |  |  |
| N Sham - E Sham | -0.0215 | 0.0505 | 957 | -0.425 | 0.9741 |
| N Sham - N Infected | 0.0253 | 0.0438 | 957 | 0.579 | 0.9385 |
| **N Sham - E Infected** | **0.1246** | **0.0438** | **957** | **2.844** | **0.0235** |
| E Sham - N Infected | 0.0468 | 0.0437 | 957 | 1.072 | 0.7069 |
| **E Sham - E Infected** | **0.1461** | **0.0437** | **957** | **3.342** | **0.0048** |
| N Infected - E Infected | 0.0993 | 0.0357 | 957 | 2.780 |  |
| *B. Female fecundity, during 96—120 hours post-infection* |  |  |  |  |  |
| **Sham N - Infected N** | **-5.10** | **1.38** | **465** | **-3.695** | **0.0014** |
| Sham N - Sham E | 1.22 | 1.34 | 465 | 0.911 | 0.7989 |
| Sham N - Infected E | -1.33 | 1.34 | 465 | -0.999 | 0.7503 |
| **Infected N - Sham E** | **6.32** | **1.38** | **465** | **4.576** | **<.0001** |
| **Infected N - Infected E** | **3.77** | **1.38** | **465** | **2.729** | **0.0333** |
| Sham E - Infected E | -2.55 | 1.34 | 465 | -1.910 | 0.2253 |

Supplementary table 1. [continued]
| Comparison | Estimate | SE | DF | t ratio | p-value |
| --- | --- | --- | --- | --- | --- |
| *C. Bacterial load upon death (BLUD)* |  |  |  |  |  |
| N Female - E Female | 0.0473 | 0.428 | 12.18 | 0.110 | 0.9995 |
| **N Female - N Male** | **1.3954** | **0.405** | **7.81** | **3.443** | **0.0367** |
| **N Female - E Male** | **1.6723** | **0.490** | **12.10** | **3.412** | **0.0229** |
| E Female - N Male | 1.3481 | 0.486 | 11.64 | 2.772 | 0.0715 |
| **E Female - E Male** | **1.6250** | **0.443** | **10.31** | **3.669** | **0.0182** |
| N Male - E Male | 0.2769 | 0.427 | 11.98 | 0.648 | 0.9141 |

**Supplementary table 2.** Pairwise comparisons using Holm-Sidak method of p-value correction for comparison between systemic bacterial loads in E_1-4_ and N_1-4_ flies at every sampling time point (hours post-infection, HPI). Significant effects (p < 0.05) are marked in bold font.

| HPI | Sum square | Mean square | F<sub>1,76</sub> value | p-value | Corrected p-value | Significance |
| --- | --- | --- | --- | --- | --- | --- |
| *A. Within-host bacterial dynamics, females* |  |  |  |  |  |  |
| **3** | **32.584** | **32.584** | **11.017** | **0.001388** | **0.020618925** | * |
| **6** | **157.07** | **157.07** | **39.621** | **1.811E-08** | **3.8031E-07** | *** |
| **9** | **297.91** | **297.91** | **46.193** | **2.117E-09** | **4.6574E-08** | *** |
| **12** | **391.04** | **391.04** | **30.513** | **4.576E-07** | **9.15196E-06** | *** |
| **15** | **283.09** | **283.09** | **13.569** | **0.0004313** | **0.006878523** | ** |
| **18** | **352.73** | **352.73** | **9.7851** | **0.002494** | **0.034355585** | * |
| **21** | **889.14** | **889.14** | **25.172** | **0.000003353** | **6.37051E-05** | *** |
| **24** | **621.56** | **621.56** | **15.266** | **0.0001942** | **0.003296276** | ** |
| 27 | 242.04 | 242.04 | 6.6574 | 0.01173 | 0.111298075 | ns |
| **30** | **281.21** | **281.21** | **9.35** | **0.003043** | **0.03884473** | * |
| 33 | 190.65 | 190.65 | 6.8927 | 0.01037 | 0.10833571 | ns |
| 36 | 65.477 | 65.477 | 4.6115 | 0.03499 | 0.220667664 | ns |
| 39 | 45.649 | 45.649 | 6.1707 | 0.01519 | 0.128691303 | ns |
| **42** | **181.41** | **181.41** | **20.215** | **0.00002479** | **0.000446126** | ** |
| 45 | 7.3113 | 7.3113 | 1.3993 | 0.2405 | 0.42315975 | ns |
| 48 | 4.8192 | 4.8192 | 1.6401 | 0.2043 | 0.681032239 | ns |
| 54 | 3.9686 | 3.9686 | 1.4655 | 0.2298 | 0.648104221 | ns |
| 60 | 0.44946 | 0.44946 | 0.1574 | 0.6927 | 0.6927 | ns |
| 66 | 20.081 | 20.081 | 6.1654 | 0.01523 | 0.115539426 | ns |
| 72 | 6.0684 | 6.0684 | 1.4543 | 0.2316 | 0.54630701 | ns |
| 84 | 13.39 | 13.39 | 4.2547 | 0.04239 | 0.228862123 | ns |
| 96 | 39.935 | 39.935 | 7.6924 | 0.006973 | 0.080540333 | ns |
| *B. Within-host bacterial dynamics, males* |  |  |  |  |  |  |
| **4** | **81.754** | **81.754** | **25.465** | **0.000002992** | **8.97597E-06** | *** |
| **10** | **347.83** | **347.83** | **50.812** | **4.018E-10** | **1.6072E-09** | *** |
| **48** | **32.364** | **32.364** | **13.025** | **0.0005482** | **0.001096099** | ** |
| 96 | 1.1459 | 1.1459 | 0.4838 | 0.4888 | 0.4888 | ns |

